# Limit of detection of *Salmonella* ser. Enteritidis using culture-based versus culture-independent diagnostic approaches

**DOI:** 10.1101/2024.02.05.578949

**Authors:** L.M. Bradford, L. Yao, C. Anastasiadis, A.L. Cooper, B. Blais, A. Deckert, R. Reid-Smith, C. Lau, M.S. Diarra, C. Carrillo, A. Wong

**Affiliations:** Department of Biology, Carleton University, Ottawa, Ontario, Canada; Research and Development, Ottawa Laboratory (Carling), Canadian Food Inspection Agency, Ottawa, Ontario, Canada; Institute for Advancing Health Through Agriculture, Texas A&M University, College Station, Dallas-Fort Worth, Texas, United States; Public Health Agency of Canada, Ottawa, Ontario, Canada; Guelph Research and Development Centre, Agriculture and Agri-Food Canada, Guelph, Ontario, Canada; Centre for Foodborne Environmental and Zoonotic Diseases, Public Health Agency of Canada, Guelph, Ontario, Canada

## Abstract

In order to prevent the spread of foodborne illnesses, the presence of pathogens in the food chain is monitored by government agencies and food producers. The culture-based methods currently employed are sensitive but time-and labour-intensive, leading to increasing interest in exploring culture-independent diagnostic tests (CIDTs) for pathogen detection. However, sensitivity and reliability of these CIDTs relative to current approaches has not been well established. To address this issue, we conducted a comparison of the limit of detection (LOD_50_) for *Salmonella* between a culture-based method and three CIDT methods: qPCR (targeting *invA* and *stn*), metabarcode (16S) sequencing, and shotgun metagenomic sequencing. Samples of chicken feed and chicken caecal contents were spiked with *Salmonella* serovar Enteritidis and subjected to culture-and DNA-based detection methods. To explore the impact of non-selective enrichment on LOD_50_, all samples underwent both immediate DNA extraction and an overnight enrichment prior to gDNA extraction. In addition to this spike-in experiment, feed and caecal samples acquired from the field were tested with culturing, qPCR, and metabarcoding. In general, LOD_50_ was comparable between qPCR and shotgun sequencing methods. Overnight microbiological enrichment resulted in an improvement in LOD_50_ with up to a three log decrease, comparable to culture-based detection. However, *Salmonella* reads were detected in some unspiked feed samples, suggesting false-positive detection of *Salmonella*. Additionally, the LOD_50_ in feeds was three logs lower than in caecal contents, underscoring the impact of background microbiota on *Salmonella* detection using all methods.

**IMPORTANCE:** The appeal of CIDTs is increased speed with lowered cost, as well as the potential to detect multiple pathogen species in a single analysis and to monitor other areas of concern such as antimicrobial resistance genes or virulence factors. Understanding the sensitivity of CIDTs relative to current approaches will help determine the feasibility of implementing these methods in pathogen surveillance programs.

## INTRODUCTION

Foodborne pathogens inflict a serious health and economic toll worldwide. In Canada, 4 million cases of foodborne illness are thought to be domestically acquired annually, with norovirus, *Clostridium perfringens*, *Campylobacter* spp, and non-typhoidal *Salmonella* the most prevalent causes of disease [1]. Detection of food pathogens throughout the food supply chain is thus critical to reduce the incidence of foodborne illness. Typically, the detection of food pathogens for surveillance and for outbreak investigation relies on isolating viable organisms using highly sensitive, culture-based methods. Since most foodborne pathogenic bacteria such as salmonellae can cause illness at very low numbers (e.g., 7 CFU) [2], methods for their detection in foods should be able to determine their presence at similarly low numbers in an analytical unit (e.g., 1-10 CFU per 25 g sample) [3]. These highly sensitive approaches are also appropriate for commodities such as feeds, where even low doses of *Salmonella* can result in poultry colonization [4]. Unfortunately, culture-based approaches can be laborious and time-consuming. For example, the time from sample collection to positive culture for *Salmonella* is up to 7 days, involving 48-72 hours of enrichment culture, and 48-72 hours of growth on selective agar followed by biochemical testing to confirm presumptive *Salmonella* colonies [3]. In recent years, there has been increasing interest in exploring culture-independent diagnostic tests (CIDTs) such as quantitative PCR (qPCR), metabarcode sequencing, and metagenome sequencing for detecting pathogens in food [5, 6, 7] and environmental samples [8, 9], and for infectious disease diagnostics in clinical settings [10, 11, 12, 13]. These methods could offer lower costs, increased speed, and the potential to detect multiple pathogens in a single analysis. In addition, metagenome sequencing can offer insights into the presence of virulence factors [14] and antimicrobial resistance genes [15]. However, pivoting to use such methods is only possible if the sensitivity and reliability of CIDTs is proven to be comparable to current approaches.

The poultry production chain is a good model for evaluating novel detection and surveillance methods, such as CIDTs. A large proportion of foodborne illnesses are associated with consumption of contaminated poultry meat [16]. In the USA, over 25% of foodborne outbreaks with known sources were attributed to poultry products [17].

Worldwide, a majority of the cases of salmonellosis and campylobacteriosis have been associated with poultry [17, 16]. Poultry products are also commonly contaminated with *Staphylococcus aureus, Listeria monocytogenes, Clostridium perfringens* and pathogenic *Escherichia coli* [18]. *Salmonella* can be introduced into poultry through feeds and persist throughout the food chain, resulting in contamination of animals and subsequent fecal contamination of retail poultry products [19, 20]. Given the importance of poultry as a protein source in the global food supply, pathogen reduction in this commodity could have important human health implications.

To address the question of whether CIDTs are adequately sensitive for detection of pathogens in food-relevant matrices, we conducted a comparison of the limit of detection (LOD_50_) for the current culture-based *Salmonella* detection method in use at the Canadian Food Inspection Agency (CFIA) and for three CIDTs (qPCR, metabarcode sequencing, and metagenomic sequencing) in samples of chicken feed and chicken caecal contents spiked with known quantities of *Salmonella*. We further assessed the use of qPCR and 16S sequencing for *Salmonella* detection in naturally contaminated caeca and feed.

## MATERIALS AND METHODS

### Caecal and feed samples

Caeca from freshly sacrificed 35 day old Ross 708 broiler chickens were from an ongoing study at Agriculture and Agri-Food Canada (Guelph, Ontario). All experimental procedures were approved (Protocol number # No. 3521) by the institutional ethics committees on animal experimentation according to guidelines of the Canadian Council on Animal Care. Samples of the broiler finisher feed which included corn as the principal cereal, and soya and soybean cake as protein concentrates (Aviagen, Huntsville, United States) were used for the feed experiments. Caeca were transported on ice and stored at 4°C overnight. Feed was stored at 4 °C until use. Starting materials were confirmed to be *Salmonella*-free by subjecting a subset to overnight incubation in buffered peptone water (BPW), DNA extraction, and marker-gene qPCR as described below.

### Overnight *Salmonella* cultures

*Salmonella enterica* ser. Enteritidis isolate CFIAFB20140150 previously isolated from raw retail poultry (accession CP133565-CP133567; Cooper et al. [21]) was used for spiking. Bacteria were revived from a glycerol stock and plated on non-selective agar. A single isolated colony was selected and inoculated into 5mL buffered peptone water (BPW; Oxoid), and incubated for 24 hr at 37 °C with 150 rpm shaking. Previous tests of overnight cultures suggested this should result in growth to 2.5*x*10^9^ CFU/mL. Overnight cultures were diluted in a 10X series in glucose-free M9 minimal medium (see supplementary methods), and these dilutions were used for spiking and for enumeration via either dropping or spreading on non-selective agar followed by overnight incubation at 37 °C. Expected vs. actual CFU spiked in are shown in Table S6 and Table S7.

### Spiking procedure

#### Caecal contents

Chicken caecal contents were “milked” into petri dishes using sterile gloves. Sterile scoops were used to transfer 0.25 g to screw-cap tubes and 1 g to pre-dispensed 9 mL falcon tubes of BPW. Screw-cap and falcon tubes containing caecal content were spiked with between 4 and 10 µL of the appropriate dilution of the *Salmonella* ser. Enteritidis culture. Spiked caecal contents in screw-cap tubes were stored at −80 °C prior to DNA extraction. For microbiological enrichment according to MFHPB-20 [3], spiked caecal contents in BPW were incubated for 21 hr at 35 °C with 100 rpm shaking.

#### Feed

For direct extractions, 10 g portions of feed were added to a filtered stomacher bag (Nasco Sampling/Whirl-Pak, United States), to which 20 mL BPW was added. The sample was homogenized using a stomacher (Interscience Laboratories, United States) for 1 minute at 230 rpm. Approximately 10 mL of liquid was recovered from each sample. Samples were subjected to a low speed spin (500 x g for 5 min) to remove eukaryotic cells.

After transfer of supernatant to a new falcon tube, samples were subjected to a high speed spin (11000 x g for 5 min) to pellet bacterial cells. Supernatants were discarded and the pellet was resuspended in 0.1 mL of BPW. The appropriate number of *Salmonella* cells were then added (Table S7).

For microbiological enrichments, 10 g portions of feed were added to a filtered stomacher bag, to which 90 mL of BPW was added. The sample was homogenized as described above, then spiked with 1 mL containing the appropriate dilution of *Salmonella* cells (Table S7). Samples were incubated for 20 hr at 37 °C.

### Growth in selective broths and agar

Recovery of *Salmonella* through secondary enrichment and growth on differential/selective agars was conducted as described in MFHPB-20 [3]. From the BPW enrichment, 1 mL was added to 9 mL of Tetrathionate Brilliant Green (TBG; Becton, Dickinson and Company, New Jersey, USA) broth and 0.1 mL to 9 mL of Rappaport-Vassiliadis Soya Peptone (RVS; Oxoid) broth. Inoculated TBG and RVS were incubated for 24 hr at 42.5 °C with 100 rpm shaking. Broths were then vortexed briefly and streaked onto Brilliant Green Sulfa (BGS; Becton, Dickinson and Company) agar and Brilliance™ *Salmonella* agar (Becton, Dickinson and Company) plates using 10 µL loops. Plates were incubated for 24 hr at 35 °C, then examined for colonies indicative of *Salmonella*.

Suspected *Salmonella* colonies were confirmed using colony PCR. For caecal content samples, colonies were picked into 100 µL TE buffer, which was heated to 100 °C for 10 minutes then cooled to 20 °C. Boiling prep material was used as a template for qPCR reactions. Reaction and temperature profiles are described in the qPCR section below. For feed samples, presumptive *Salmonella* colonies were confirmed by PCR amplification of the *invA* gene (Table S1). Each 25 µL reaction contained 1x GoTaq Colourless Master Mix (Promega, United States) and 0.3 µM Primers (invA_1869F, invA_1999R). Colony material was transferred directly into the PCR mix, and was patched onto brain-heart infusion agar. PCR cycling conditions were as follows: denaturation at 95°C for 2 min, followed by 40 cycles of 95 °C for 30 s, 60 °C for 30 s, 72 °C for 30 s, followed by a final extension at 72°C for 5 min. PCR products were visualized by capillary electrophoresis using a QIAxcel DNA high-resolution gel cartridge on a QIAxcel instrument (Qiagen, Toronto, Canada), according to manufacturer’s instructions.

### DNA extraction

DNA extraction was performed using the DNeasy PowerSoil Pro Kit (Qiagen, Toronto, Canada) according to kit protocols. For extraction from enriched caecal samples, the remaining volume of BPW enrichments were centrifuged at 500 xg for 5 min to pellet solids. Two mL of supernatant were centrifuged at 14000 xg for 5 min and the cell pellet was transferred to a PowerBead pro tube. For directly extracted samples, frozen spiked caecal content was thawed and beads from a PowerBead pro tube were added to the screw-cap tubes. For enriched feed samples, 10 mL of enrichments were centrifuged at 500 xg for 5 min to pellet solids. The supernatant was transferred to a new tube and centrifuged at 14000 xg for 5 min and the cell pellet was transferred to a PowerBead pro tube. For direct extractions from feed, the spiked cell pellets were transferred to PowerBead pro tubes. DNA was eluted in 100 µL of elution buffer and quantified with PicoGreen (Thermo Fisher Scientific, Canada) according to the manufacturers’ recommendations.

### Detection of marker genes by quantitative PCR

Detection of *Salmonella* based on the presence of marker genes *invA* and *stn* was performed by multiplex qPCR. Each reaction contained 12.5 µL of Roche FastStart Essential DNA Probes Master (Sigma-Aldrich, Oakville, Canada), 0.4 µM each *invA* primers, 0.3 µM each *stn* primers, 0.2 µM each probe (Table S1), 2.5 µL of DNA template, and water to a total volume of 50 µL. Cycling conditions are given in Table S2. The DNA template per reaction was 935 ng for caecal content samples, 3.75 ng for enriched feed samples, and 24 ng for unenriched feed samples. DNA concentrations were chosen based on standardization to the lowest sample concentration within a given group, and DNA input for enriched feed samples was further diluted to prevent overloaded reactions. Non-template controls received 2.5 µL PCR-grade water instead of DNA template. qPCR reactions were performed in triplicate. Duplicate standard curves in 10X dilution series from 106 to 1 genome copies per µL were run on each qPCR plate. The qPCR was performed on a Bio-Rad CFX Opus 96 Real-Time PCR System (Bio-Rad Laboratories Ltd., Mississauga, Canada) using the following temperature program: 95 °C for 5 min, followed by 45 cycles of 95 °C denaturing for 10 s, 58 °C annealing for 15 s, 72 °C extension for 10 s, and a final cooling step of 37 °C for 30 s. Two different cycle thresholds were established for determining positivity for Samonella: 40 cycles, based on the lack of any amplifications in no-template controls, and a more stringent setting of 35 cycles as is commonly used in food safety monitoring programs.

### Sequencing

The 16S-V4 and shotgun sequencing was performed at the McGill Genome Centre, and 16S-V3-V4 sequencing was performed at the CFIA Ottawa (Carling) laboratory. Samples were selected for 16S and shotgun sequencing based on results of culture-dependent and qPCR tests (Table S6 and Table S7).

Primers 16S-F_515F and 16S-R_806R (Table S1, Caporaso et al. [22]) were used to amplify the 16S V4 variable region in PCR reactions using Kapa HiFi Hotstart ready mix (Sigma-Aldrich, Oakville, Canada) (Tables S2, S3). Amplicon sequencing libraries were prepared according to the 16S Metagenomic Sequencing Library Preparation protocol [23] and sequenced with PE150 on an Illumina NovaSeq6000.

Primers 16S-F_341F and 16S-R_785R (Table S1, Klindworth et al. [24]) were used to amplify the 16S V3-V4 variable region (Tables S2, S3). Amplicon sequencing libraries were prepared according to the 16S Metagenomic Sequencing Library Preparation protocol [23] and sequenced with PE300 on an Illumina MiSeq.

Shotgun sequencing libraries were prepared using the Lucigen NxSeq AmpFREE Low DNA Library Kit (VWR International, Radnor, USA), and sequenced with PE150 on an Illumina NovaSeq6000.

### Bioinformatic analysis

#### 16S

Analysis of 16S sequence data (both V4 and V3-V4 regions) was performed in QIIME2 v2022.11 [25]. Primers were removed with cutadapt using anchored forward and reverse sequences, with –p-match-read-wildcards –p-match-adapter-wildcards to account for variations in degenerate primer sequences. Untrimmed reads were discarded. Trimmed reads were denoised with DADA2 [26]. V4 amplicons were denoised with no truncation then merged with a minimum overlap of 4 nt. Representative reads were classified using the q2-feature-classifier plugin [27] and the pre-trained Naive Bayes classifier silva-138-99-515-806-nb-classifier.qza [28] [29], available from the QIIME2 data resources site. V3-V4 amplicons were denoised with truncation at base 260 on the forward read and 190 on the reverse read, then merged with a minimum overlap of 12 nt. Representative reads were classified using the q2-feature-classifier plugin and a Naive Bayes classifier trained on the 341-785 region of the silva 138 database [29]. Following classification, mitochondria and chloroplast ASVs were removed using the filter-table plugin. QIIME2 output files were imported into R 4.2.3 [30] using the qiime2r package [31] and results were visualized using the phyloseq package [32].

#### Shotgun

Shotgun sequencing datasets were analyzed according to the pipeline established in Bradford et al. [33]. Custom workflows were made in snakemake [34]. Briefly, reads were trimmed and quality-selected with Trimmomatic [35] using the parameters minlength 36, sliding window 4:20. All passing reads, whether paired and unpaired (forward or reverse), were retained for the best chance of *Salmonella* detection. For caecal content samples, host reads were removed by classifying passing reads with Kraken 2 [36] against a custom-made Kraken 2 database made using the *Gallus gallus* reference genome from NCBI (GRCg6a; GenBank accession GCA_000002315.5). For feed samples, reads were classified against the Kraken 2 plant database. Details on these databases can be found in the supplementary material. Reads matching the host database were removed using the filterbyname function of BBMAP [37], producing quality-controlled, host-free datasets. These reads were then classified using Kraken 2, with confidence set at 0.25, using a bacteria database downloaded using the kraken2-build command on Oct 28, 2021. All reads classified as members of the *Salmonella* genus were extracted using the filterbyname function of BBMAP. The blastx function from the Blast suite [38, 39] was used to compare putative *Salmonella* reads against a blast-formatted database of *Salmonella* “species”-specific regions from [40]. Samples with reads that were called as *Salmonella* by Kraken 2 and then passed this confirmation step are considered to be positive for *Salmonella*.

Reads in the unspiked (negative control) feed samples which were identified as *Salmonella*-derived via this pipeline were tested against the NCBI-nt database via the web interface. Megablast was used with default settings, excluding results from *Salmonella* (taxid:590), using the nt database posted on April 23, 2023.

### Enrichment broth dilution test

It is possible that the carrying capacity of BPW was quickly reached in caecal spiking experiments due to the high bacterial load. This would limit the possible number of divisions of *Salmonella* spiked into the broth. To determine if dilution of the caecal contents can decrease the LOD_50_ of *Salmonella*, a dilution series was conducted using 10 additional caeca obtained from Agriculture and Agri-Food Canada (Guelph, Ontario). Contents from 10 caeca were mixed and split amongst 16 tubes (Fig. S4). Tubes were spiked with 0 (unspiked control), 3.5, 35, or 3.5*x*10^6^ (positive control) CFU of *Salmonella* enterica ser. Enteritidis isolate CFIAFB20140150 grown in BPW, as above. Each tube was then diluted 1:10 until the 10^3^ dilution was reached (Figure S4). After an overnight incubation, DNA was extracted using the DNeasy PowerSoil Pro Kit (Qiagen, Toronto, Canada) according to kit protocols, as above. Detection of *Salmonella* based on the presence of marker gene *invA* was performed as described above.

### Limit of detection calculations

LOD_50_ of each method and condition combination was calculated according to Wilrich and Wilrich [41] using the tool provided at https://www.wiwiss.fu-berlin.de/fachbereich/vwl/iso/ehemalige/wilrich/index.html.

### Plotting and statistical analyses

Plotting and statistical analyses were performed in R v4.2.3 [30]. A full list of packages used can be found in the Supplementary Methods (subsection R packages).

### Proof of concept experiment

Feed and chicken caeca were sent to labs at the CFIA and the Public Health Agency of Canada (PHAC) for *Salmonella* testing as part of their ongoing monitoring programs. These samples underwent culture-based detection following the MFHPB-20 protocol, and aliquots of the non-selectively-enriched material were provided to us for DNA extraction and testing via CIDTs. DNA extraction, multiplex qPCR, and sequencing of the V3-V4 regions of the 16S rRNA gene were performed as described above. In total, 56 caeca samples and 48 feed samples were tested.

## RESULTS

We compared the limit of detection (LOD_50_) of enrichment-culture based *Salmonella* detection methodology against three culture-independent diagnostic tests (CIDTs): qPCR, 16S sequencing, and metagenomic sequencing. We spiked two matrix types (chicken caecal contents and chicken feed) with known quantities of *S.* Enteriditis. For the CIDTs, all samples underwent both immediate DNA extraction and an overnight enrichment incubation in non-selective media to investigate the impact of this enrichment step.

### Detection is strongly influenced by matrix

Across all methods and enrichment conditions, *Salmonella* could be detected at much lower spike-in levels in feed samples, which have low microbial abundance, than in caecal contents. The lowest LOD_50_ in feed samples was 0.047 CFU/g (via culturing), compared to 50 CFU/g for caecal contents (via post-enrichment qPCR) (Fig. **1**). *Salmonella* could not be detected in caecal contents via 16S sequencing, regardless of enrichment condition.

**FIG 1.**
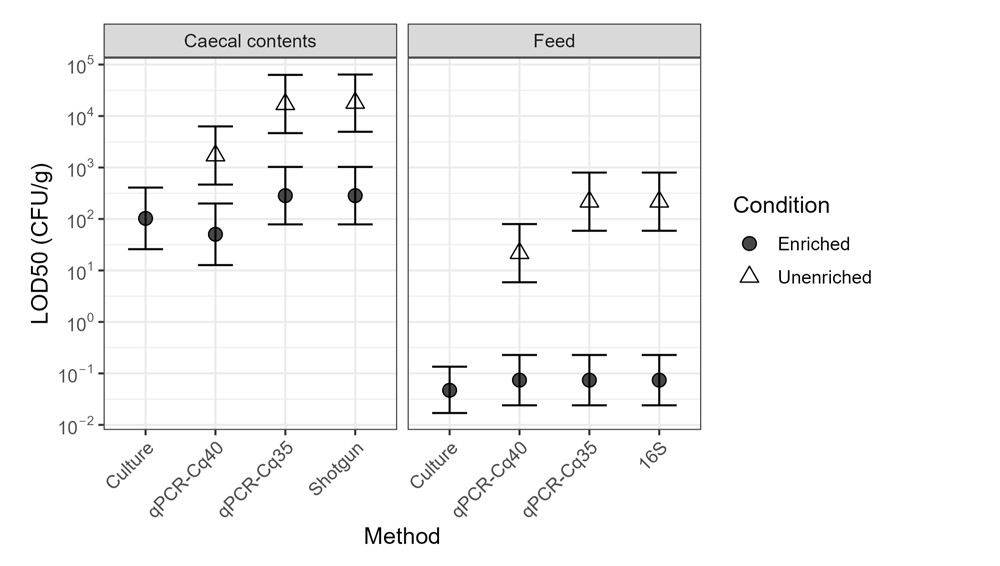
Limits of detection for the methods and conditions tested according to the log-log model by Wilrich and Wilrich [41]. Note that no *Salmonella* was detected in caecal contents by 16S sequencing, and LOD_50_ could not be calculated for shotgun sequencing analysis of feed samples because all samples were positive. Calculations assume no *Salmonella* was detected in negative controls. 16S represents V3-V4 amplicon sequencing. qPCR-Cq40 and -Cq35 represent qPCR with Cq cutoffs of 40 and 35 cycles, respectively. Error bars show 95 % confidence intervals.

### Enrichment enhances detection

In the absence of enrichment, CIDTs had considerably worse LOD_50_ than traditional, culture-based testing (Fig. **1**). In caeca, shotgun metagenomics and qPCR with a Cq cutoff of 40 had LOD_50_ approximately 2-log higher than culture-based detection; use of a Cq cutoff of 35 provided an improvement of 1-log. Lack of sensitivity was more pronounced in feed, where 16S, qPCR-35, and metagenomics had LOD_50_ 3-log higher than culturing.

The first step of the culture-dependent method is an overnight incubation in BPW. In order to evaluate the impact of this initial incubation on test sensitivity, DNA was extracted directly from spiked samples (“unenriched”) and from the BPW post-incubation (“enriched”), and these DNA extracts were used for CIDTs. The majority of reads within the shotgun sequencing datasets from unenriched feed came from plants, reducing the usable data; in contrast, plant-derived reads were a tiny proportion in the enriched feed datasets (Fig. S3). Although BPW is not selective for *Salmonella*, enrichment lowered the LOD_50_ in all methods in which both conditions were tested. The LOD_50_ of CIDTs using DNA extracted directly from caecal contents was particularly high, at 1.7*x*10^3^ CFU/g for qPCR (40 cycle threshold) and 1.8*x*10^4^ CFU/g for metagenomics via shotgun sequencing. With enrichment, the LOD_50_ of these methods dropped to 50 and 283 CFU/g, respectively. The effect was even more pronounced in feed samples, where, for example, LOD_50_ of qPCR was 21.7 CFU/g without enrichment but 0.074 CFU/g with enrichment (Fig. **1**).

Enrichment was performed with 9 mL of BPW to 1 g of material as described in the culture-detection protocol [3]. Diluting caecal contents to raise the BPW:material ratio improved detection, as shown with qPCR-based detection of the *invA* gene (Fig. S4). Of the six replicate samples spiked with 10 CFU/g *Salmonella* in this dilution experiment, *invA* could be detected in just one at the 9:1 ratio, in three replicates after a 10X dilution, and in all six replicates after a 100X dilution (Fig. S4).

### Enrichment has varying effects on community composition

Sequencing of 16S rRNA shows that overnight enrichment in BPW had a noticeable effect on the community composition of feed samples (Figs. **2**, S1). The Enterobacteriaceae family, to which *Salmonella* belongs, was only a small proportion of the community prior to enrichment but rose to >50 % post-enrichment, concurrent with a drop in alpha diversity (Fig. S2). Multiple genera within the Enterobacteriaceae greatly increased their proportion of the community during enrichment, including potentially pathogenic *Citrobacter*, *Klebsiella*, *Escherichia-Shigella*, and *Salmonella* (Fig. **2**). The Actinobacter and Bacilli classes decreased in abundance, and Clostridia sequences appeared in a few feed samples following enrichment. Conversely, the overall community composition in caecal content samples showed little change (Fig. S1), and diversity dropped only slightly in enriched vs. unenriched samples (Fig. S2). Enterobacteriaceae were <= 2.5 % of the unenriched community and rose to 5-13 % of communities post-enrichment, but the majority of Enterobacteriaceae sequences belonged to the *Escherichia-Shigella* genus, as defined by the Silva v138.1 database [29]. Sequences representing *Salmonella* were not found in any of the caecal samples selected for 16S sequencing. The most abundant class in the caecal contents was Clostridia, which comprised 89-97 % of unenriched and 77-93 % of enriched caecal communities (Fig. S1). Clostridia families Lachnospiraceae and Ruminococcaeceae were 3-28 % and 4-14 % of the total communities, respectively.

**FIG 2.**
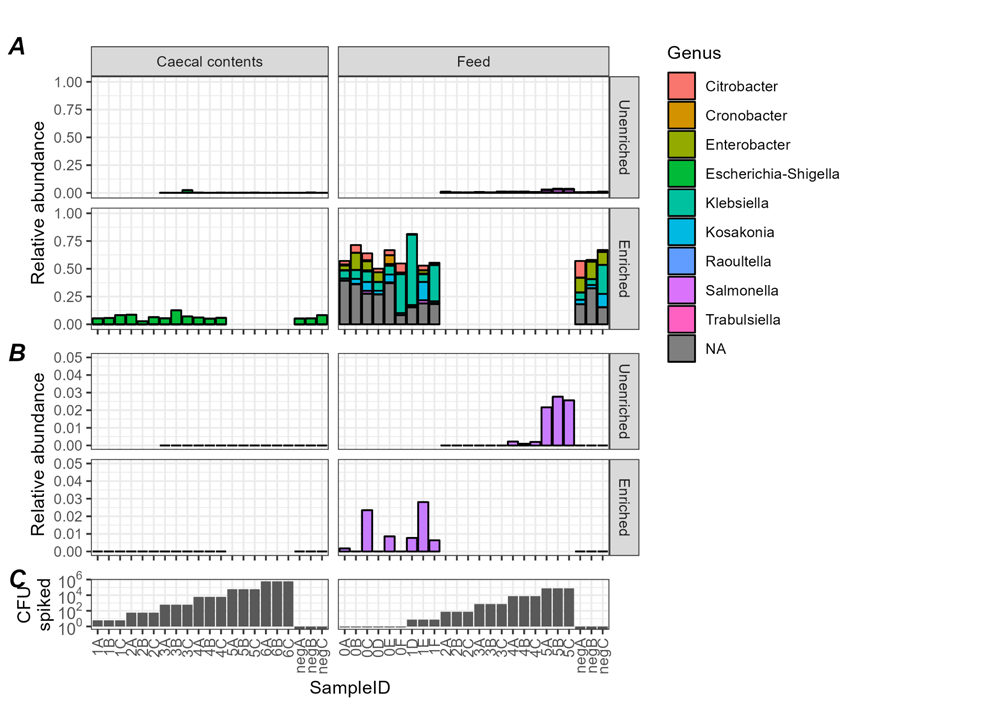
(A) Relative abundance of genera in the Enterobacteriaceae family according to sequencing of the 16S V3-V4 region. Colours indicate assigned genus, with “NA” indicating sequences that could not be assigned below the family level. (B) Zoomed-in view showing only the *Salmonella* genus abundance from V3-V4 sequencing. Note the scale of the y-axis. Blank areas are shown for samples that were not sequenced. (C) Number of *Salmonella* Entertidis CFU spiked into samples in above panels.

### Possible false positives for *Salmonella* in feed

Evidence of *Salmonella* was not found in unspiked feed samples via culturing or 16S rRNA analysis. However, one gene targeted by the multiplex qPCR (*invA*) amplified with high Cq values in two of the three enriched unspiked feed samples (Fig. **3**). According to the draft protocol, amplification of either target indicates that a sample may be positive for *Salmonella*. All three enriched unspiked feed samples, as well as two of three unenriched unspiked feed samples, were found to contain shotgun sequencing reads classified as *Salmonella*-derived according to our analytical pipeline (Table S8). We carried out further investigations to determine whether these samples were in fact contaminated with *Salmonella*, or if they represent false positives. We were able to isolate and sequence colonies of *Citrobacter* species from additional feed samples and found that some sequencing reads were considered to have come from *Salmonella* when tested with our shotgun sequencing pipeline (see Supplementary Methods). These *Citrobacter* isolates, however, do not contain the *invA* gene that is tested for with qPCR.

**FIG 3.**
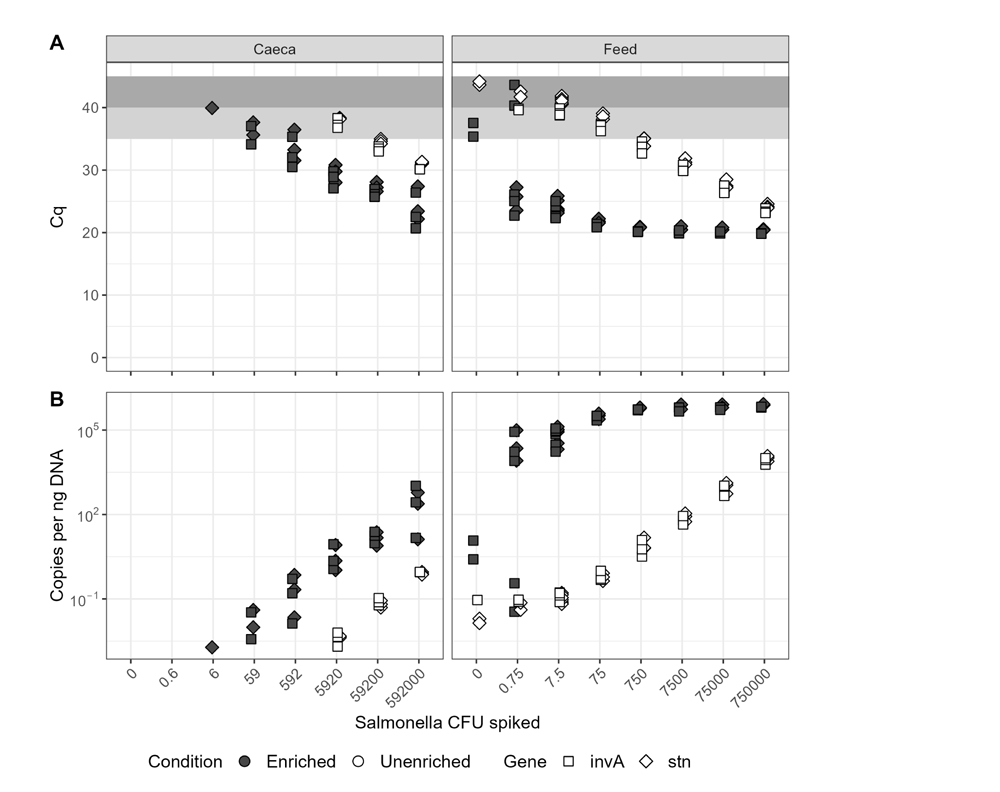
Detection of *Salmonella* marker genes via multiplex quantitative PCR. (A) Cq values. Samples with data points in the dark grey zone above 40 cycles are considered negative; samples with data points in the light grey zone between 35 and 40 cycles may be interpreted as positive; samples with data points below 35 are definitely positive. (B) Gene copies per ng of input DNA, as calculated using standard curves. Y-axis is in log scale.

### qPCR-based detection is comparable to culturing in naturally contaminated samples

Following the spike-in experiments, a proof-of-concept experiment was performed on chicken feed and caecal contents acquired by the Canadian Food Inspection Agency and the Public Health Agency of Canada as part of their food safety monitoring programs. Culture-based testing was performed by these government agencies, and the post-enrichment material was sent to us for DNA extraction and testing by CIDTs. There was very strong concordance between detection by culturing and by multiplex qPCR. When qPCR positivity is defined as Cq values < 40, detection results were identical. If qPCR positivity is set more stringently with a 35 Cq threshold, 14 of the 19 culture-positive samples were found to be positive by qPCR. Detection via sequencing of the 16S rRNA V3-V4 regions was much less sensitive in these samples, with only one feed and two caecal samples determined to be positive for *Salmonella* via this method (Fig. **4**). The samples found to be positive by 16S sequencing had low Cq values in the multiplex qPCR assay (Fig. **4**).

**FIG 4.**
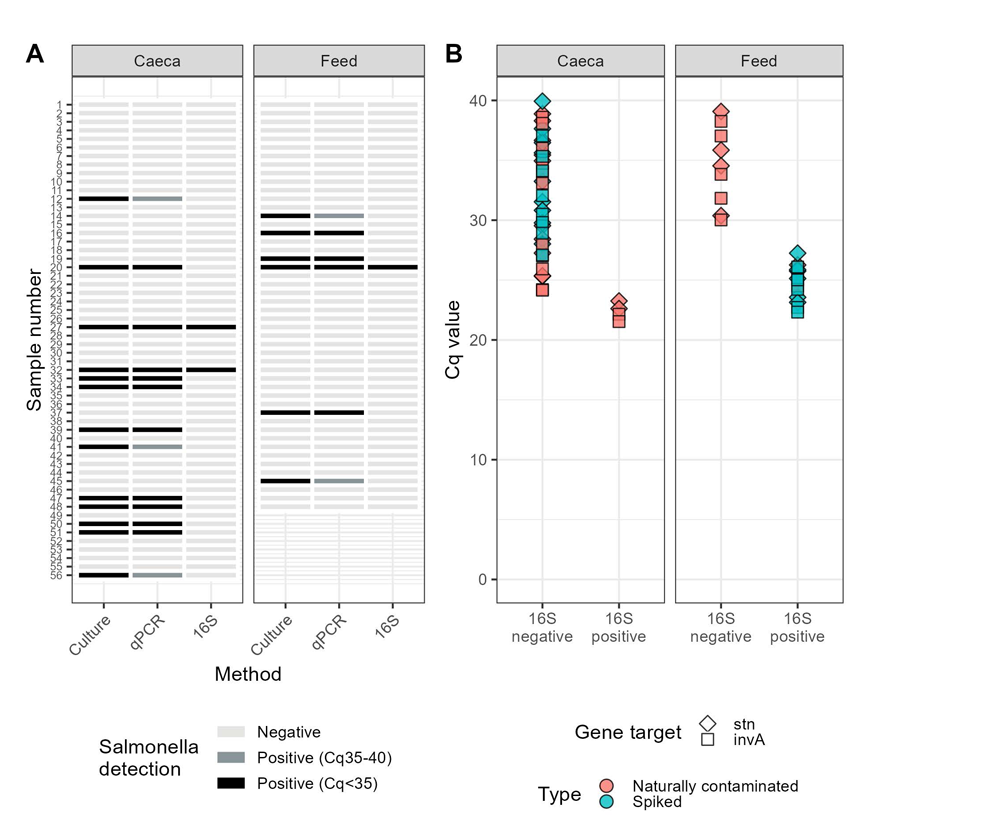
(A) Results of *Salmonella* detection by culturing and CIDTs on enriched natural samples. (B) Comparison of Cq values from qPCR against positive/negative detection of *Salmonella* via sequencing of the 16S V3-V4 variable regions. Separate Cq values are plotted for the two gene targets in the multiplex qPCR assay. Results shown are from enriched samples which showed amplification in qPCR reactions and which underwent 16S sequencing.

## DISCUSSION

The primary question driving this investigation was whether various CIDTs have sufficient sensitivity and reliability to be used in food safety applications. To answer this, we systematically compared limits of detection (LOD_50_) for current enrichment-culture based methodology against three culture-independent diagnostic tests (CIDTs). We focused on *Salmonella* as a model pathogen, and on matrices relevant to poultry production: chicken feed (low bacterial load) and chicken caecal contents (high bacterial load). Within each matrix, we found the LOD_50_ for CIDTs to be equivalent to that of the culture-dependent method when using DNA from material that underwent an overnight enrichment in non-selective broth (Fig. **1**). Testing DNA extracted directly from *Salmonella*-spiked matrices yielded a higher LOD_50_ in every case. Although enrichment is time-consuming, it is essential for detection sensitivity using CIDTs, as has been found for *Salmonella* [42, 43] and other bacterial pathogens in food matrices [44].

There was good concordance between detection via culturing and multiplex qPCR on enriched materials, as has been seen with various qPCR methods [45, 46, 47, 48]. Although culturing, qPCR, and sequencing of the 16S rRNA V3-V4 region had equivalent LODs when tested on spiked enriched feed samples, only qPCR was able to match culturing results when used on naturally contaminated samples. Sequencing depth and quality were well-matched between these two investigations. Samples in which 16S sequencing could detect *Salmonella* were those with lower Cq values in qPCR analysis, indicating that a higher proportion of *Salmonella* DNA within the samples was needed for 16S detection with the method and sequencing depth we used. Reduced relative proportions of *Salmonella* in enrichment cultures derived from naturally-contaminated samples are likely indicative of an extended lag time for *Salmonella* growth, attributed to damage to the organism due to environmental stress conditions. Additionally, only one strain of *Salmonella* ser. Enteritidis was used in the spike-in portion of this study. Different strains and serovars may have variable growth kinetics in enrichment culture [49].

Metabarcoding was undertaken using both the V4 and V3-V4 variable regions of the 16S rRNA gene. Amplicons of the V4 region are an appropriate length (approx. 291 nt) for sequencing on Illumina HiSeq or NovaSeq, which can produce millions of reads per sample, providing comprehensive information on bacterial community compositions. However, *Salmonella* sequences in this region are not unique and reads cannot be distinguished between *Salmonella* and other genera within the Enterobacteriaceae [50]. Amplicons of the V3-V4 region are longer (approx. 464 nt), and can be used to specifically detect *Salmonella*. The read length required was best suited to an Illumina MiSeq, yielding much lower read depths per sample. Though targeted read depths per sample exhibit significant variation among different studies, 100 million reads represent a reasonable quantity, and it is unlikely that laboratories engaged in routine monitoring would surpass this threshold. Some of the positive results obtained from the *Salmonella* qPCR assays had Cq values that were higher than 35 cycles. Interpretation of high Cq values may be complicated as these may represent false-positive results [51]. High Cq values could be generated by degradation of probes, contamination, or by non-specific amplification of nucleic acids present in complex samples. In a diagnostic lab, enrichments that were qPCR positive, but with high Cq values may be further investigated by increasing the amount of sample (e.g. gDNA) loaded, or by trying to recover target organisms, but these results on their own would not be conclusive. In this study we observed “true positives” with Cq values of 40 cycles, however, some of the unspiked feed samples had a signal at this threshold. Ultimately, further evaluation of the method is needed to empirically determine reliable Cq cutoffs in a variety of matrices. In our study we tried to maximize the amount of the gDNA sample loaded in the PCR assay to increase the relative proportion of the sample being used in the assay, particularly for the direct extraction from spiked samples. Genomic DNA from the samples was eluted into 100 µL of liquid, therefore each qPCR assay included about 2.5 % of the total sample. Total gDNA extracted from caecal contents was much higher than for feed, resulting in use of almost 1 ug gDNA/assay for caecal samples. Further dilution to normalize feed and caecal concentrations would have significantly decreased the proportion of the sample loaded in the assay, which would have consequently impacted LOD_50_.

All methods had very low LOD_50_ (0.047 – 0.074 CFU/g) in enriched feed samples, although unenriched LOD_50_ varied. This can likely be attributed to the fact that *Salmonella* cells spiked into feed were unstressed and readily viable, having just been grown in an overnight culture in rich broth. Other microorganisms on the feed had, conversely, been subsisting on dry feed material at cool (4 °C) temperatures. The goal of non-selective enrichment is to allow recovery of stressed or injured cells, but it is easy to imagine that healthy *Salmonella* enjoyed a competitive advantage over the feed microbiome in this environment, thus artificially decreasing post-enrichment LODs. For this study we elected to forgo the stressing procedures that would typically be used in a method validation study to avoid complications associated with variability introduced by this procedure. The LOD_50_ for stressed cells would likely be somewhat higher than observed here. Caecal contents, on the other hand, were freshly harvested from chickens and processed after a single night of storage at 4 °C, thus minimizing stress on the resident microbiota. The majority of the caecal content community, both with and without enrichment, belongs to the Clostridia class (Fig. S1), which are common constituents of the gastrointestinal tracts of omnivorous, warm-blooded animals [52]. The abundance of members of this class is consistent with surveys of chicken caecal communities [52]. All Clostridia are obligate anaerobes [53], which would not be expected to maintain an overwhelming presence after enrichment in an oxic environment. One possible explanation is that, due to the high biomass in caecal content, the carrying capacity of the broth was quickly reached with very little opportunity for growth of aerobes. Results of an experiment in which caecal contents were serially diluted in BPW before overnight enrichment support this hypothesis, with improved qPCR-based detection in samples with higher BPW:caecal content ratios during enrichment (Fig. S4).

The relatively high LOD_50_ for *Salmonella* in caecal contents have implications for monitoring schemes that rely on testing these materials, notably the National Microbiological Baseline Study in Broiler Chicken December 2012 [54]. That study suspended chicken caecal contents in a 1:4 (w/w) ratio with BPW, then screened using the BAX PCR system (Hygenia, Mississauga, Canada), with presumptive positives enumerated by Most Probably Number (MPN) culturing. They found that 25.6 % of the caecal samples tested were positive for *Salmonella*, with 65 % of those positives enumerated at > 110 MPN/g. However, our results suggest that the positivity rate may have been higher, but hidden by the inability of *Salmonella* to grow sufficiently during enrichment. Our findings may also have implications for other studies and monitoring schemes that test for pathogens in high biomass backgrounds such as probiotic preparations and fermented consumable products [55, 56].

While buffered peptone water (BPW) is considered a non-selective medium, we found clear evidence that overnight growth in BPW favours the growth of some taxa to the exclusion of others. Non-selective enrichment of feed caused profound changes in the bacterial community compositions. Previous studies on non-selective enrichment (using BPW or Universal Pre-enrichment Broth, UPB) of various food products saw a decrease in proportion of Proteobacteria (which includes *Salmonella*) and an increase in Firmicutes, with varying results for Actinobacteriota [57, 58, 59]. Conversely, non-selective enrichment in our experiment caused an increased proportion of Proteobacteria, decrease in Firmicutes, and the near-disappearance of the Actinobacteriota phylum. The Proteobacteria phylum consisted mostly of members of the Enterobacteriaceae family, including *Citrobacter*, *Klebsiella*, *Escherichia-Shigella*, and *Salmonella* genera. There is thus a need for further work on the effects of enrichment on the microbial communities of different commodities.

Amplification of the *invA* gene during qPCR and detection of putatively *Salmonella*-derived shotgun sequencing reads in unspiked feed sample controls suggest that *Salmonella* DNA may have been present. This does not guarantee the presence of viable cells; indeed, the inability of CIDTs to distinguish between viable cells and lingering DNA is a known downfall of these methods [60, 61]. It is also possible that signals were generated from nonspecific products generated in these complex samples [62]. The number of reads identified as coming from *Salmonella* was higher in enriched samples than in their unenriched counterparts, which could indicate growth of viable cells. The more likely explanation is that these reads are false positives due to presence of related organisms. We previously isolated a *Citrobacter werkmanii* from the feed used in this experiment which contains sequences matching those found in the unspiked feed controls, and have since isolated multiple *Citrobacter* colonies from feed with sequences that are attributed to *Salmonella* in our bioinformatic pipeline. Characterization of these isolates is ongoing. *Citrobacter* spp. are closely related to *Salmonella* [63] and have been shown to cause false positives during food testing [64, 65]. The genome of the previously isolated *Citrobacter* has not been uploaded to NCBI or other databases, so it was not available during determination of the *Salmonella* species-specific regions used during bioinformatic analysis [40], although shotgun reads simulated from its genome were tested during pipeline development and did not result in false *Salmonella* hits [33]. Read classification in metagenomic analysis relies on matching sequences to curated databases [66]. Over-representation of pathogenic species in public repositories relative to commensal organisms commonly found in food and environmental species has the potential to lead to false-positive detection of pathogens as observed in this study [67]. This emphasizes the need for caution when using CIDTs for food safety or in health diagnostics.

CIDTs are promising tools for pathogen surveillance and detection in agriculture, food safety, and medicine. However, the performance of CIDTs must be systematically investigated to guide their appropriate use. Here, we show that the CIDTs tested have equivalent sensitivity to culture-based detection methods when an overnight incubation is employed, but much higher limits of detection (that is, lower sensitivity) without this enrichment. Detection limits of all methods are clearly influenced by the matrix background, which must be considered when interpreting results from varied matrices. We also show the major downside of CIDTs, i.e., the potential for false positives and lack of cultured isolates on which to perform further tests.

## ACKNOWLEDGMENTS

Thank you to the personnel at the Public Health Agency of Canada and the Canadia Food Inspection Agency who provided samples for the proof of concept portion of this research.

## DATA AVAILABILITY STATEMENT

The data have been deposited to NCBI with links to BioProject accession number PRJNA1035945. Code can be found at https://github.com/LMBradford/SalmLOD-paper

## CLINICAL TRIALS

Not applicable.

## ETHICS APPROVAL

All experimental procedures were approved (Protocol number # No. 3521) by the institutional ethics committees on animal experimentation according to guidelines of the Canadian Council on Animal Care.

## Supplemental material

A supplementary file containing supplementary methods, figures, and tables is provided as a separate PDF.

